# High arsenic levels increase activity rather than diversity or abundance of arsenic metabolism genes in paddy soils

**DOI:** 10.1101/2021.05.03.442544

**Authors:** Si-Yu Zhang, Xiao Xiao, Song-Can Chen, Yong-Guan Zhu, Guo-Xin Sun, Konstantinos T. Konstantinidis

## Abstract

Arsenic (As) metabolism genes are generally present in soils but their diversity, relative abundance, and transcriptional activity in response to different As concentrations remain unclear, limiting our understanding of the microbial activities that control the fate of an important environmental pollutant. To address this issue, we applied metagenomics and metatranscriptomics to paddy soils showing a gradient of As concentrations to investigate As resistance genes (*ars*) including *arsR*, *acr3*, *arsB*, *arsC*, *arsM*, *arsI*, *arsP*, and *arsH* as well as energy-generating As respiratory oxidation (*aioA*) and reduction (*arrA*) genes. Somewhat unexpectedly, the relative DNA abundances and diversity of *ars*, *aioA*, and *arrA* genes were not significantly different between low and high (∼10 vs ∼100 mg kg^-1^) As soils. By comparison to available metagenomes from other soils, geographic distance rather than As levels drove the different compositions of microbial communities. Arsenic significantly increased *ars* genes abundance only when its concentration was higher than 410 mg kg^-1^. In contrast, between low and high As soils, metatranscriptomics revealed a significant increase in transcription of *ars* and *aioA* genes, which are induced by arsenite, the dominant As species in paddy soils, but not *arrA* genes, which are induced by arsenate. These patterns appeared to be community-wide as opposed to taxon-specific. Collectively, our findings advance understanding of how microbes respond to high As levels and the diversity of As metabolism genes in paddy soils and indicated that future studies of As metabolism in soil, or other environments, should include the function (transcriptome) level.

**IMPORTANCE:** Arsenic (As) is a toxic metalloid pervasively present in the environment. Microorganisms have evolved the capacity to metabolize As, and As metabolism genes are ubiquitously present in the environment even in the absence of high concentrations of As. However, these previous studies were carried out at the DNA level and thus, the activity of the As metabolism genes detected remains essentially speculative. Here, we show that the high As levels in paddy soils increased the transcriptional activity rather than the relative DNA abundance and diversity of As metabolism genes. These findings advance our understanding of how microbes respond to and cope with high As levels, and have implications for better monitoring and managing an important toxic metalloid in agricultural soils and possibly other ecosystems.

## INTRODUCTION

Arsenic (As) is a toxic metalloid, widely distributed in Earth’s crust. Relatively high amounts of As are introduced into the environment from nature sources such as volcanic eruptions, weathering of rocks and geothermal activities as well as anthropogenic activities such as mining, pigment production and application of As-based pesticides in agriculture (1, 2). Because of its pervasive presence in the environment, microorganisms have evolved the capacity to metabolize As (3), and this capacity is hypothesized to be an ancient mechanism that emerged at least 2.72 billion years ago (4, 5).

Microbial mediated As biotransformation include As detoxification to mitigate toxicity and As respiration to generate energy. Several mechanisms have evolved to detoxify both inorganic and organic As. The inorganic arsenate [As(V)] compounds in cytoplasm can be reduced by a reductase (ArsC) (6), followed by arsenite [As(III)] efflux outside the cell, which is catalyzed by two evolutionarily unrelated As(III) efflux permeases, i.e. ArsB and Acr3 (7). Additionally, As(III) can be methylated by an As(III) *S*-adenosylmethionine (SAM) methyltransferase (ArsM) to the more toxic organic methylarsenite [MMAs(III)] (8), followed by the organoarsenical detoxification pathways, which have been recently shown to be present in various microbial genomes. The latter detoxification pathways include MMAs(III)’s oxidization to methylarsenate [MMAs(V)], which is catalyzed by the MMAs(III)-specific oxidase ArsH (9), and actively transported outside the cell by the MMAs(III) efflux permease ArsP (10), or MMAs(III)’s demethylation to the less toxic As(III) by the ArsI lyase, which cleaves the carbon-As bond in MMAs(III) (11). These As resistance genes are usually organized in an *ars* operon, which is controlled by one of three known ArsR transcriptional repressors (ArsR1, ArsR2 and ArsR3) regulated selectively by As(III) and one (ArsR4) by MMAs(III), depending on the exact version of the operon (12, 13). The As respiratory pathways include oxidation of As(III) either to detoxify it to less toxic As(V), or couple it to ATP production (14), and dissimilatory reduction that couples As(V) reduction to anaerobic heterotrophic growth (15). As(III) oxidation is catalyzed by As(III) oxidase (Aio) whose large subunit (AioA) has been well characterized in both bacteria and archaea (16). On the other hand, As(V) reduction is catalyzed by respiratory As(V) reductase (Arr), whose large catalytic subunit (ArrA) serves as a reliable marker for the process (17).

Due to the anaerobic conditions typically prevailing in paddy soils, As(III) is commonly the predominant form of As, which is more mobile and bioavailable to microbes for biotransformation (18–20). Indeed, microbial genes involved in As resistance such as *arsC* and *arsM*, and genes involved in As respiratory such as *aioA* and *arrA* have been detected to be widely presented in the paddy soils with As contamination (21), and their activity to biotransform As has also been experimentally demonstrated (22–24). Recent studies of soils with low As levels, i.e. typically less than 15 mg kg^-1^ (or ppm), also showed relative high abundance and diversity of microbial As metabolism genes; most notably, *arsB*, *acr3*, *arsH*, *arsR, arsC*, *arsM*, *aioA* and *arrA* genes in paddy soils (25) and *arsC*, *arsM*, *aioA* and *arrA* genes in estuarine sediments across southeastern China (26). These results reveal high prevalence of As metabolism microbes in the environment even in absence of high concentration of As. However, these previous studies were carried out at the DNA level and thus, the expression of the As metabolism genes detected remains speculative. Further, these DNA-based results showing prevalent As metabolism microbes in both high and low As level environments are somewhat surprising because gene functions and species selected by a specific factor tend to be more abundant in ecosystems where the factor (selective pressure) is stronger as this has been shown for various soil factors such as pH, total organic carbon (TOC), total nitrogen (TN), phosphorus (TP) (27) and organic pollutants (28).

Herein, we applied both metagenomics and metatranscriptomics to investigate the resident (at the DNA level) and active (at the RNA level) As-associated microbial genes in paddy soils of different As concentrations, aiming to address the following questions: (1) Do As metabolism genes in high As levels paddy soils show higher relative abundances and/or transcriptional activities than their counterparts in low As levels paddy soils to cope with the higher As level present? (2) Which are the key As metabolism genes and pathways coping with high As levels? (3) What are other factors than As levels driving microbial communities compositional and gene transcriptional differences in soils with As contamination?

## RESULTS

### Soil physicochemical characteristics

A total of 18 samples, three biological replicates from the same field for each site, were collected from six paddy soils in Hunan, China with different As concentrations (Fig. S1 and Table S1). Arsenic (As) concentrations in these paddy soils varied significantly (ANOVA, *p* < 0.01) between high (67.3-104.0 mg kg^-1^) and low (2.5-10.8 mg kg^-1^) As levels. The high As concentrations in the paddy soils likely resulted from the historic mine activities occurring in the same area. The total concentrations of nitrogen (TN) in high As paddy soils were also significant higher (ANOVA, *p* < 0.01) than those with low As (1.64-1.77 g kg^-1^ vs 1.17-1.24 g kg^-1^). No significant difference was found in the other soil physiochemical characteristics, including pH, Eh, total concentration of carbon (TC), phosphorus (TP), sulfur (TS), and concentrations of heavy metals including Pb, Cd, Cr, Sb and Fe. Details of the soil properties were summarized in supplementary Table S2.

### Broad microbial community diversity indices

Six shotgun metagenomes and six shotgun metatranscriptomes were obtained after the triplicate DNA and RNA samples from each site were pooled together, respectively. For metagenomes (Table S3), 12-15 Gbp (billion base pairs, 13 Gbp, on average) of shotgun metagenomic data and an average read length of ∼115 bp were acquired after trimming for each pooled DNA sample. On average, 6.6 Gbp (ranged from 3.9 to 8.3 Gbp) of shotgun metatranscriptomic data and average read length of ∼129 bp were obtained after trimming and removal of rRNA reads for each pooled RNA sample (Table S4). The coverage achieved by sequencing based on the read redundancy value estimated by Nonpareil was around 22-27%, except for sample SKS_AsHig, which had a lower alpha-diversity (measured by Nonpareil) than the others (*Nd* 22.6 vs 23.6-23.8; note that *Nd* is in log scale) and thus, higher coverage (42%). No significant difference (*p* > 0.05) was found in the alpha-diversity between high and low As paddy soils (average of 23.3 vs 23.7). The average genome size of microbial community was 5.7 Mbp and 5.6 Mbp in high and low As paddy soils, respectively. Considering the difference of average genome size of microbial community between these paddy soils was no more than 2-fold, estimation of As metabolism genes abundance was essentially the same between the RPKG and RPKM metrics.

### Paddy soil community taxonomic composition

16S rRNA gene (16S) fragments extracted from the metagenomes were about 0.03-0.04% of the total reads (Table S3) and around 65-70% of them were classifiable at the phylum level. Bacteria were predominant in these paddy soils, accounting for 61-69%, while Archaea accounted for only 0.5-2.4%; the remaining reads were unclassified against the SILVA database (29). The dominant bacteria comprised of phyla Proteobacteria, Acidobacteria, Chloroflexi, Nitrospirae, Verrucomicrobia and Bacteroidetes, representing 50-59% of the total 16S-carrying reads. Archaea community was mostly represented by the Crenarchaeota phylum at 0.05-0.78% of the total, followed by Euryarchaeota at 0.36-0.77% (Fig. 1A). There were overall no significant relationships between the level of As and community composition, assessed by Bray-Curtis similarity in 16S rRNA gene-based OTUs (NMDS plot; Fig. S2). DESeq2 analysis (Fig. 1B) revealed statistically significant lower abundance of members of Euryarchaeota, i.e. *Methanomicrobia* and *Thermoplasmata* (0.15% vs 0.85% and 0.02% vs 0.07%, respectively; *p* < 0.05) in high vs low As paddy soils. In contrast, the abundance of Chloroflexi, mostly comprised of Ellin6529 (1.44% vs 0.52%), S085 (0.28% vs 0.09%), *Thermomicrobia* (0.06% vs 0.01%), and *Chlorobi*-OPB56 (0.16% vs 0.05%), GAL15 (0.21% vs 0.01%) and OP11-WCHB1-64 (0.07% vs 0.01%) were significantly (*p* < 0.05) higher in high vs low As paddy soils. We also attempted to recover metagenome-assembled genomes (MAGs; see details in supplementary Results and Discussions), which yielded only five MAGs of high quality (completeness >80% and contamination <5%) presumably due to the high complexity of the datasets. These MAGs were assignable to the class of *Deltaproteobacteria*, *Thermomicrobia*, *Clostridia*, *Acidobacteriaceae* and *Solibacteres*, which is consistent with the dominant microbes in these sites as revealed by the read-based findings reported above.

**Figure 1.**
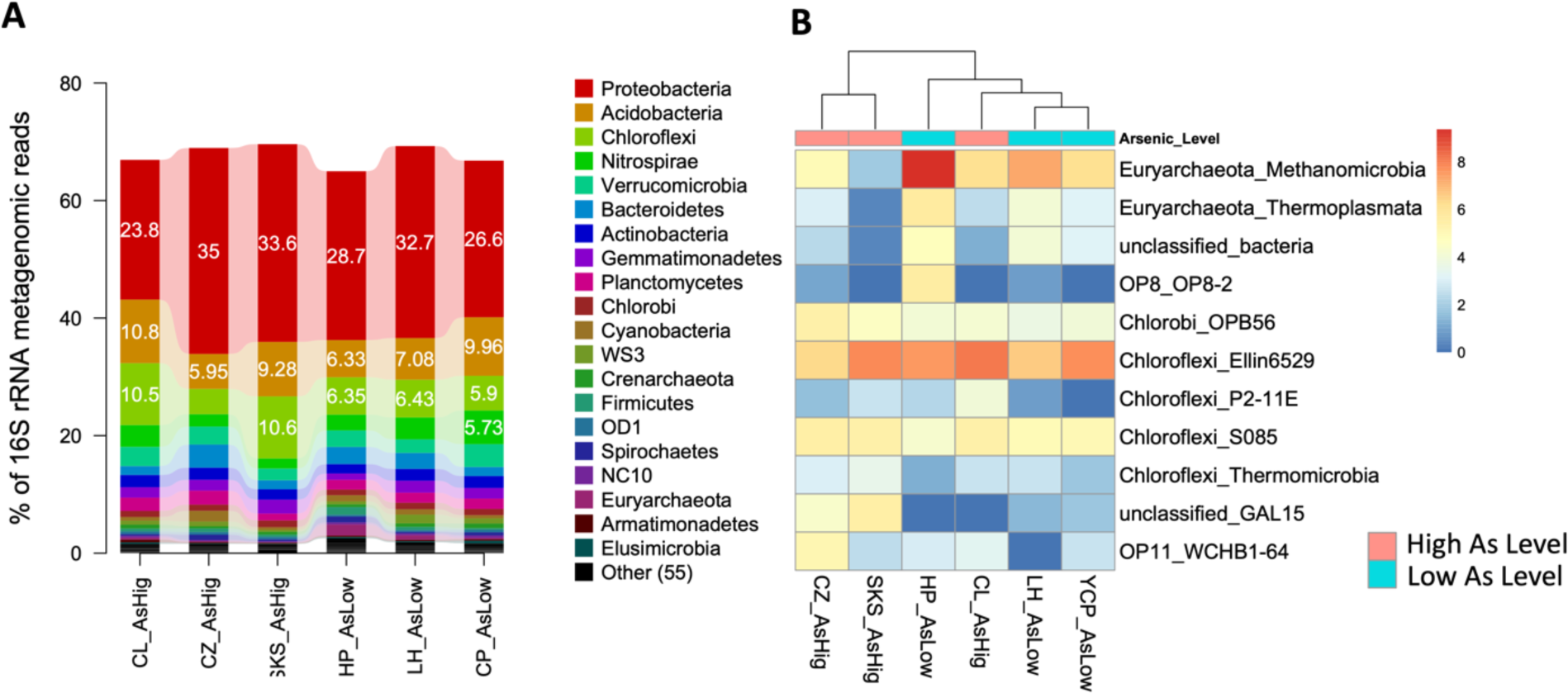
Comparison of microbial community composition in high vs low As sites. (A) Phylum-level community composition and abundance of microbes in paddy soils based on taxonomic classification of identified 16S rRNA gene sequence fragments. The relative abundance of each taxon was normalized by the total number of 16S rRNA gene sequence fragments (both assigned and unassigned) obtained in each corresponding metagenome. Only the top 20 most abundant phyla are shown. (B) Heatmap prokaryotic classes showing statistically significant differences in abundance between high (red) and low (blue) As paddy soils (negative binomial test, adjusted *p* value < 0.05).

### Paddy soil community functional composition

Protein-coding fragment sequences were predicted for about 90-92% and 44-59% of the trimmed reads in metagenomics and metatranscriptomics, respectively (Table S3 and S4). The high As microbial communities clustered separately than their low As counterparts based on functional gene content (summarized as annotation counts from SEED subsystems) albeit they were also quite different from each other and thus, did not cluster tightly together (Fig. 2A). It was not unexpected to observe clustering at the functional but not taxonomic composition of the sampled communities considering that functional gene content has been shown to be more strongly and faster influenced by environmental conditions than phylogenetic beta-diversity (30). DESeq2 analysis revealed no statistically significant changes in the relative abundance of genes predicted from metagenomics of different As levels. In contrast, genes associated with As resistance, phage tail fiber, phage photosynthesis, phage capsid, phage cyanophage and CBSS-316279 were all significantly (adjusted *p* < 0.05) more expressed based on metatranscriptomes by a log2 fold change of 1-2.3 in high vs low As paddy soils (Fig. 2B and 2C).

**Figure 2.**
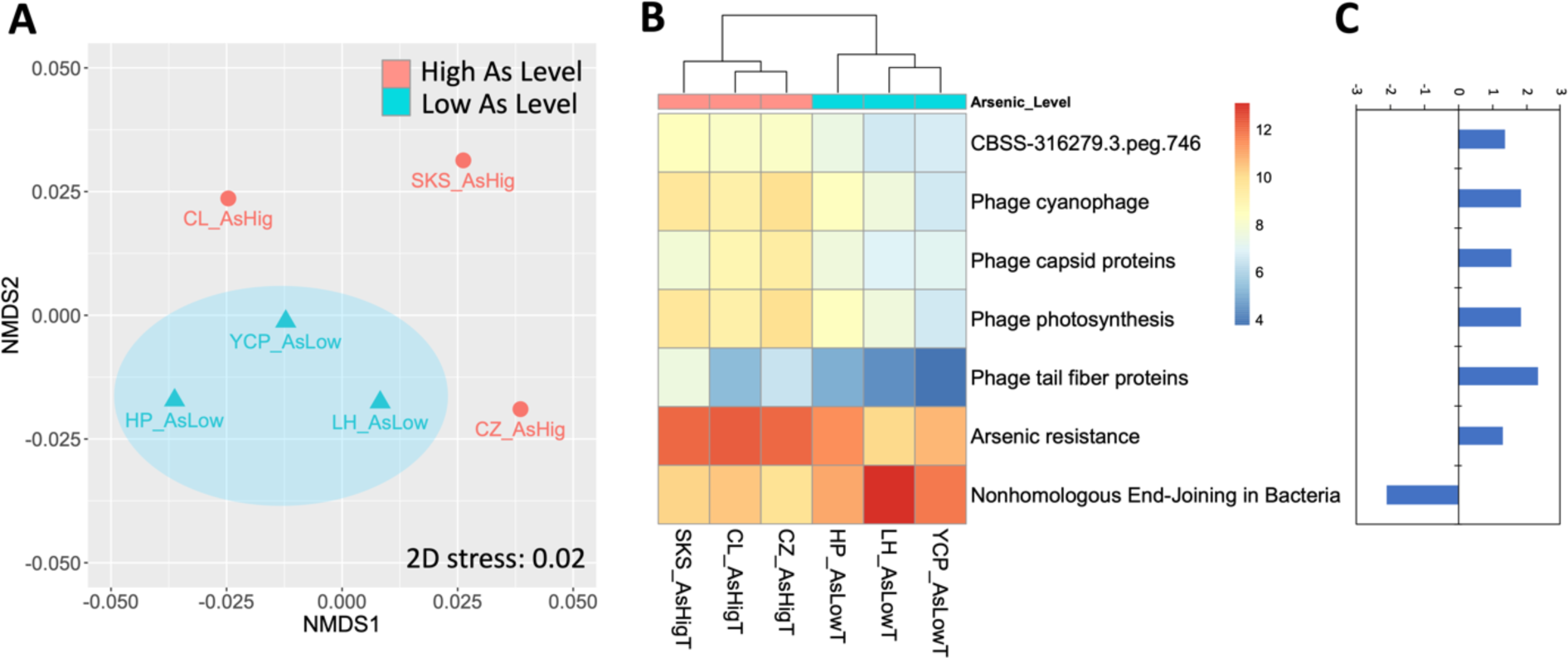
The effect of high As levels on the functional composition of microbial communities. (A) NMDS plot of the Bray-Curtis metric based on the counts of SEED subsystems (level 3) annotated from metagenomes in high (red) and low (blue) As paddy soils. (B) Heatmap of SEED subsystems (level 1) annotated from metatranscriptomes showing statistically significant differences in abundance between high (red) and low (blue) As paddy soils (negative binomial test, adjusted *p* value < 0.05). (C) Log2 fold change of SEED subsystems (level 1) annotated from metatranscriptomes that were significantly differentially abundant between high and low As paddy soils (negative binomial test, adjusted *p* value < 0.05).

### Relative DNA abundance and expression of As resistance and respiratory genes

Overall, the relative abundance of *ars* genes, i.e., the cumulative abundance of *arsR* (*arsR1*, *arsR2*, *arsR3*, *arsR4*), *acr3*, *arsB*, *arsC* (*arsC1*, *arsC2* and *acr2*), and *arsM*, *arsI*, *arsP* and *arsH* genes, were comparable between high and low As paddy soils as revealed by metagenomes (1.2 × 10^-2^ ± 9.7 × 10^-3^ vs 6.1 × 10^-3^ ± 5.3 × 10^-4^ RPKM; Fig. 3A). An outlier to this result was sample SKS_AsHig that had three times higher relative abundance of *ars* genes compared to other samples (Fig. S3). To robustly assess transcriptome activity while accounting for DNA abundance, the transcriptional level of *ars* genes were estimated by normalizing metatranscriptomic counts by the relative abundance of same *ars* genes identified in metagenomes. More than 80% of the As metabolism genes detected in metatranscriptomes were identified in the metagenomes, representing the majority of the As metabolism genes in the samples. The normalized transcriptional expression levels of *ars* genes were significantly higher (*p* < 0.05) in high vs low As paddy soils (2074 ± 701 vs 710 ± 337 RPKM; Fig. 3A). Overall, the 13 *ars* genes responsible for As transcriptional repressor (*arsR*), As(III) efflux (*acr3* and *arsB*), As(V) reduction (*arsC* including *arsC1*, *arsC2* and *acr2*), As(III) methylation (*arsM*), MMAs(III) demethylation (*arsI*), MMAs(III) oxidation (*arsH*) and MMAs(III) efflux (*arsP*) all consistently showed higher transcriptional activities in high vs low As paddy soils (Fig. 3B). However, for sample HP_AsLow, the *arsH*, *arsR* and *arsP* showed a slightly higher transcriptional activity than one of the samples from high As levels sites (CZ_AsHig or SKS_AsHig), which could be attributed to high soil heterogeneity or biases during library preparation and/or sequencing.

**Figure 3.**
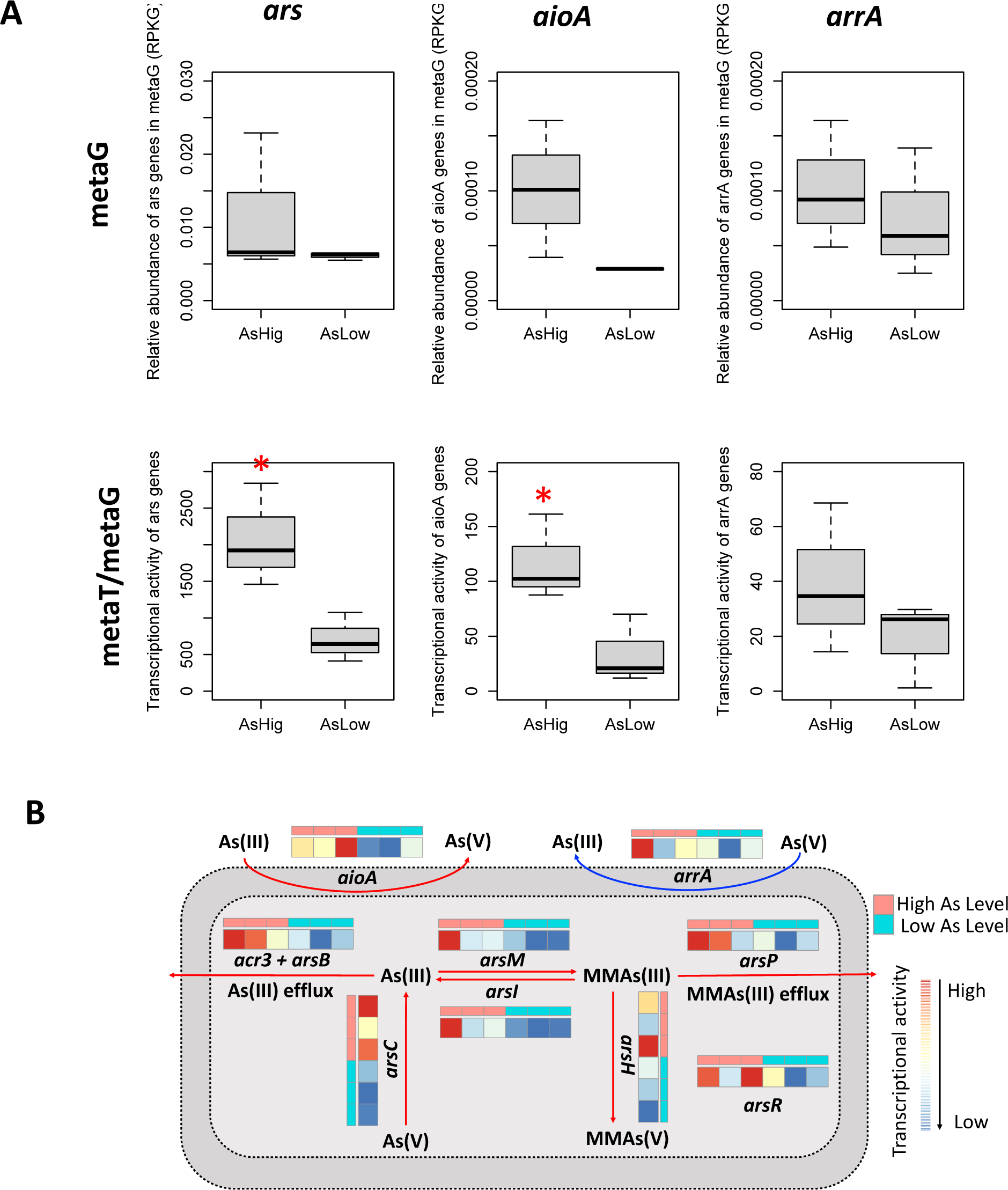
Relative abundance and transcription of *ars*, *aioA* and *arrA* genes in high and low As paddy soils. (A) The relative abundance and transcription of *ars* genes is the cumulative of *arsR* (*arsR1*, *arsR2*, *arsR3*, *arsR4*), *acr3*, *arsB*, *arsC* (*arsC1*, *arsC2* and *acr2*), *arsM*, *arsI*, *arsP* and *arsH* genes. Relative abundance of As metabolism genes in metagenomes was normalized by the statistic RPKG (reads per kb per genome equivalent, see materials and methods for more details). The As metabolism genes transcriptional activity was evaluated by the relative abundance of As metabolism genes in metatranscriptomics (RPKM reads per kb per millions of reads) normalized by the relative abundance of the same gene identified in metagenomics (RPKM). Horizontal lines represent the median value; * indicates statistically different means at *p* < 0.05 based on one-way ANOVA analysis. (B) Transcriptional activity of the As metabolism genes and the cellular location of their encoded proteins. Transcriptional activities (the same values as in Figure 3A) of *ars* (*acr3*, *arsB*, *arsC*, *arsM*, *arsI*, *arsH*, *arsP* and *arsR*), *aioA* and *arrA* genes in high (red) and low (blue) As paddy soils visualized in a heatmap that was produced using the pheatmap package in R 3.5.1. The color ranges from red to blue indicate transcriptional activity from high to low.

For *aioA* genes (Fig. 3A), which encode respiratory As(III) oxidase in the periplasm, a higher relative DNA abundance was revealed, although not statistically significant (*p* = 0.114), in high vs low As paddy soils (1.0 × 10^-4^ ± 6.2 × 10^-5^ vs 2.9 × 10^-5^ ± 3.2 × 10^-7^ PRKG, respectively). The transcriptional activity of *aioA* genes was strongly significantly (*p* < 0.05) increased in high vs low As paddy soils (117 ± 39 vs 34 ± 31 PRKM, respectively). Further, there was no significantly higher abundance (*p* = 0.598) or transcriptional activity of *arrA* genes (*p* = 0.33), which is responsible for respiratory As(V) reduction, between high and low As paddy soils (Fig. 3A). The relative abundance of *arrA* genes in metagenomes were comparable between high and low As paddy soils (1.0 × 10^-4^ ± 5.8 × 10^-5^ vs 7.4 × 10^-5^ ± 5.9 × 10^-5^ RPKG, respectively), while their transcriptional activity was slightly higher, but not significant (*p* = 0.33), in paddy soils with high vs low As levels (39 ± 27 vs 19 ± 16 RPKM, respectively).

### Responses to high As level in paddy soils are community-wide rather than taxon-specific

To evaluate which clades expressed higher As genes in high As soils, and thus, assess whether the transcriptomic shifts of As metabolism gene described above were due to systematic community-wide response as opposed to a few taxa, the Ars, AioA and ArrA-carrying reads in metagenomes (DNA reads) and metatranscriptomes (RNA reads) were placed on the their representative phylogenetic clade (typically >95% nucleotide identity within- vs <95% identity between-clades) and the ratio of RNA/DNA reads assigned at the same reference clade (or gene-based OTU) was compared between samples (Fig. 4 and Fig. S4-S12). Overall, in high As soils, more than 60% (ranging between 63% and 98%; Table S6) of the total clades recruited reads from metatranscriptomes for each As metabolism gene, except for *arsB*, suggesting that the high As levels mostly induced a community-wide response. Moreover, around 54-78% of the total clades for Ars, AioA and ArrA proteins showed a higher ratio of RNA/DNA reads in high than low As paddy soils (Table S6), revealing that the As metabolism genes were expressed more strongly in high than low As samples by many distinct clades (taxa). For *arsB*, due to relative lower abundance of this gene compared to the others (e.g. 75-375 times lower compared to *acr3* which is also responsible for As(III) efflux), only 17% of the total clades recruited *arsB*-carrying reads from high As paddy soils and no clade recruited reads from the low As paddy soils.

**Figure 4.**
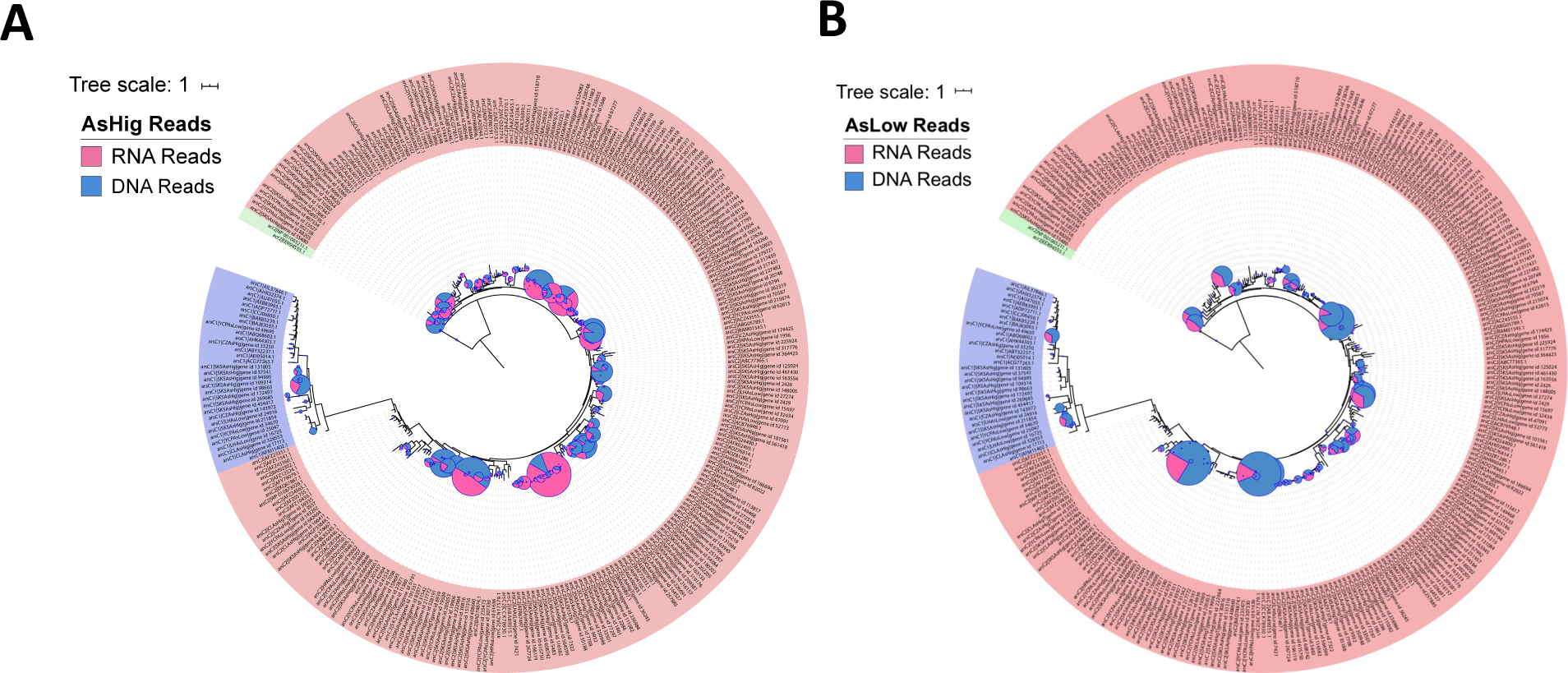
Community-wide shifts as an effect of high As levels. Phylogenetic placement of *arsC*-carrying reads identified in metagenomes (DNA reads, in blue) and metatranscriptomes (RNA reads, in red) from high (A) and low As paddy soils (B). For this analysis, the three metagenomes and metatranscriptomes from the same As level (i.e. high or low) sites were combined together. The pie size indicates the DNA abundance and the ratio indicate transcriptional activity for each clade (i.e., fraction of RNA vs DNA reads assigned to the class with the same thresholds for a match; see Methods for further details). The clades are colored on the outside based on the ArsC type, i.e., ArsC1 (blue), ArsC2 (red) and ACR2 (green). Phylogenetic placement of other As metabolism genes carrying reads are provided in the supplementary. Note that several clades in the high As recruited more RNA than DNA reads (red color dominates) whereas the opposite pattern was observed for most clades in the low As datasets.

### Environmental factors affecting the composition of microbial communities in metagenomes

To investigate the differences between microbial communities in high vs low As soils, five previously characterized samples with high As (total As concentration ranged from 34 to 821 mg kg^-1^) affected by a mine field in Hunan, China (31) and eight samples with low As (total As concentration ranged from 2 to 16 mg kg^-1^) in Hunan, Zhejiang, Jiangxi and Guangdong, China (25) were included in comparisons with the samples determined by our study. Because these metagenomes had more than two-fold variation in sequencing effort applied, we subsampled the metagenomes to the same level (2 Gbp) in order to avoid false positive signal in detecting features as differentially abundant due to differences in coverage alone (32). PCoA-based analysis of Mash distances of whole metagenomes showed that samples were mostly separated based on their geographic distances, e.g., the most distinct metagenomes were those that originated form the most distant sites (Fig. 5A). The Adonis dissimilarity test corroborated that geographic distribution patterns imposed significant (*p* < 0.01) changes in microbial community compositions. Notably, As level did not cause any significant changes (*p* > 0.05) of the whole microbial communities based on metagenomic (DNA) abundances. For the As-associated microbial communities (Fig. S13), PCoA-based analysis of Mash distances of all As gene-carrying reads recovered from metagenomes revealed a weaker effect of geographic distance in shaping diversity pattens compared to the whole microbial communities. In contrast, As level showed a relatively higher effect, especially on the As-associated microbial communities in the mine field, which is characterized by higher As level, ranging from 34.1 to 821.2 mg kg ^-1^. However, when the analysis was restricted to the paddy soils characterized by our study, no clear significant effect of high vs low As level ((∼10 vs ∼100 mg kg^-1^) was observed, which was consistent with the NMDS plot of Bray-Curtis metric based on the counts of As genes annotated from metagenomes (Fig. S14).

**Figure 5.**
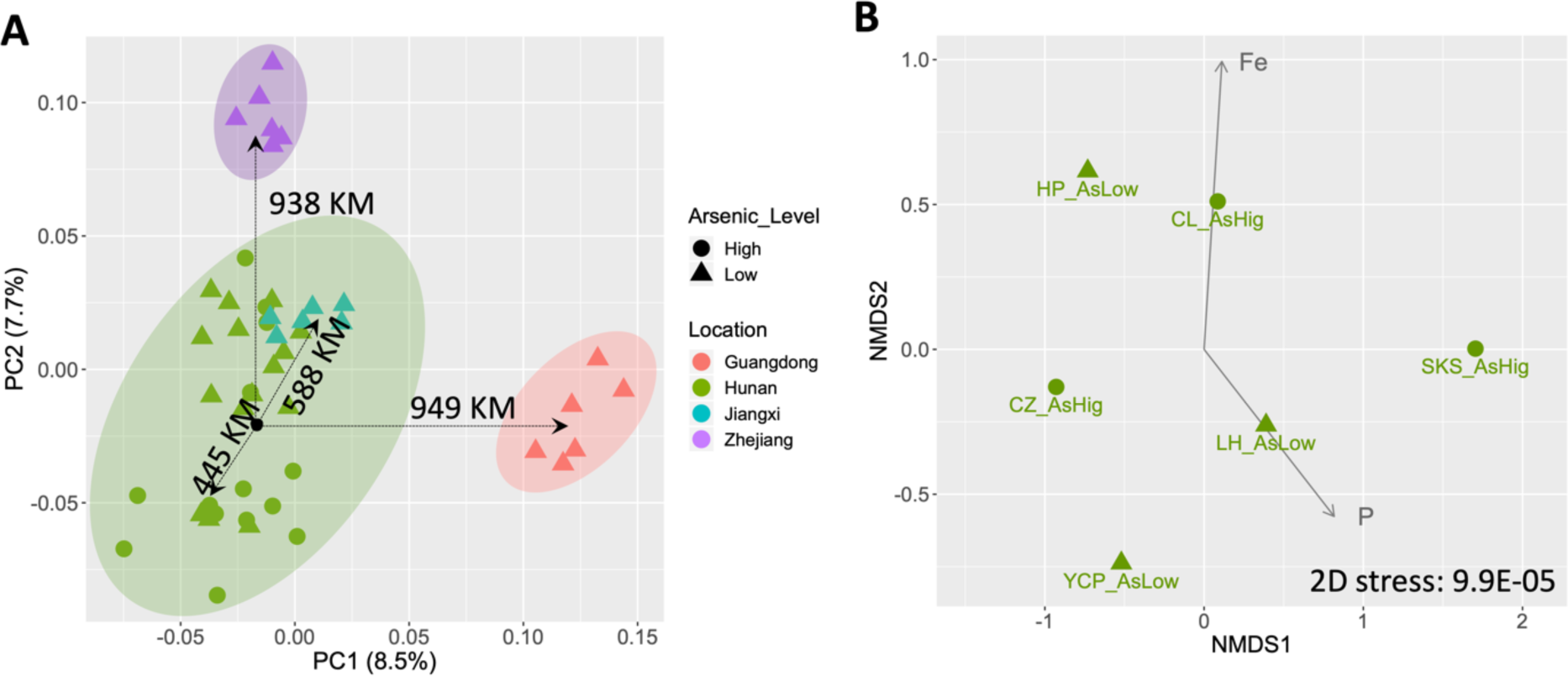
Physicochemical factors affecting whole microbial community composition. (A) PCoA plot of site-to-site similarity matrix based on Mash distances of whole metagenomes from this study and 13 selected metagenomes from previous studies. Symbols in different colors indicate samples from different provinces (green: Hunan; red: Guangdong; blue: Jiangxi; purple: Zhejiang). The shapes of symbols indicate different As levels of these samples (circle: high As levels; triangle: low As levels). Note that samples were clustered according to their geographic distance, not As levels (i.e. CL-CZ: 445 km, CL-JX: 588 km, CL-ZJ: 938 km, CL-GD: 949 km; most distinct metagenomes were clustered further apart). (B) NMDS plot of Mash distances of whole metagenomes determined by this study. The correlation of microbial community differences with environmental factors are shown (arrows). Fe and P concentrations were significantly correlated with the variations observed between the microbial communities based on the Adonis test (*p* < 0.05).

Based on the samples determined by our study, the concentrations of total phosphorus (TP) and iron (Fe), in contrast to As, were significant factors in driving the variations observed between the microbial communities (Fig. 5B), explaining about 30% of the differences among the sampled total microbial communities. A significant (*p* < 0.01) interaction was also found between these two factors. Interestingly, the TP concentration also significantly correlated with the relative abundance of different *ars* genes in the metagenomic datasets (R^2^ = 0.83, *p* < 0.05). The As concentration was found to significantly correlate with the variance in *ars* gene transcriptional activities assessed by Bray-Curtis distance between paddy soils (Adonis test, R^2^ = 0.69, *p* < 0.05) and the transcriptional activities of *aioA* genes (Pearson’s correlation = 0.85, *p* < 0.05). In addition to As, the concentration of total nitrogen (TN) (R^2^= 0.73, *p* < 0.01) and chromium (R^2^= 0.72, *p* < 0.05) also significantly correlated with differences in the transcriptional activities of *ars* genes, and the concentration of antimony (Sb) (Pearson’s correlation = 0.83, *p* < 0.05) significantly correlated with the transcriptional activities of *aioA* genes (Table S7). No significantly interaction was found between these environmental factors as revealed by PERMANOVA analysis. For the transcriptional activities of *arrA* genes, no environmental factor was found to affect these activities among the different paddy soils examined.

## DISCUSSION

Arsenic metabolism genes were pervasively present in the paddy soils either with high (67.3-105 mg kg^-1^) or low (2.5-10.8 mg kg^-1^) As levels, and their relative DNA abundance was mostly comparable between these soils (Fig. 3). Microbial capacity of As metabolism has been shown to be an ancient trait developed in response to the pervasive presence of As in early Earth (3). Therefore, it is likely that the presence and relative abundances of As metabolism genes in the paddy soils are possibly a legacy effect of an early life characteristic, at least to some extent (4). Further, the As resistant mechanisms, including the most well studied detoxification As(V) reduction system (ArsC) followed by the efflux of reduced As(III) (ACR3 and ArsB) (1), and the recently identified organoarsenical detoxification systems (ArsM, ArsI, ArsH and ArsP) (10, 33) have been detected in various microbes (34). The respiratory reduction of As(V) or oxidation of As(III), which is thought to have been developed in the Archaean era (16, 35), have also been described in a broad diversity of microbes (36). Consistently, the prevalent As metabolism genes have been previously reported in various wetlands with low As levels, such as paddy soils and estuarine sediments (< 15 mg kg^-1^) (21, 25, 26), corroborating the wide prevalence As metabolism genes in our paddy soils.

Notably, the As concentration in high As paddy soils, up to ∼100 mg kg ^-1^ in the paddy soils studied here, did not significantly correlated to a higher relative abundance of As metabolism genes in the resident As-associated microbial community. Previous study of As metabolism genes in mining impacted fields showed a significant (*p* < 0.05) linear correlation of *arsC* and *aioA* gene abundance with the increasing of As concentration from 34 to 821 mg kg ^-1^ (31). We studied the correlation of *ars* genes abundances with As concentrations using our own and the previously determined samples by Luo and colleagues and found, consistent with the previous study, a significant correlation (*p* < 0.05) between As metabolism gene abundance and high As concentration (especially in the 410-812 mg kg ^-1^ range; Fig. S15). These findings indicated that the 12 times fold higher As concentration in high (average of 92 mg kg ^-1^) vs low (average of 8 mg kg ^-1^) paddy soils of our study was not strong enough to significantly differentiate As genes abundances, and that higher difference in As level and/or high absolute As concentrations (e.g., > 410 mg kg ^-1^) is probably necessary in order to elicit more clear gene content and abundance differences. The possibilities that some of the As we measured in our high As soils is not bioavailable (and thus, not selecting for more As-metabolism genes), especially considering that the water soluble or exchangeable (bioavailable) As only accounts for about 0.4-24% of the total As in paddy soils (37). Therefore, to further corroborate these findings, however, the selection pressure exercised by different As levels, As species should also be measured in future studies.

Nevertheless, the higher abundance of *ars* (*arsR*, *acr3*, *arsB*, *arsC*, *arsM*, *arsI*, *arsP* and *arsH*) and *aioA* genes at the expression (but not DNA) level presumably reflected a truly stronger response of As genes to increased levels of As in the paddy soils at the RNA level than DNA level. Among the As metabolism proteins, both the ArsR and AioA proteins are induced by As(III), which is the dominant As species in the paddy soils during the flooded period (18, 19), and most likely contributed to the significantly increased transcriptional activities of *ars* and *aioA* genes in soils with higher As levels (Fig. 3). But the overall transcriptional activity of *arrA* gene, which is responsible for dissimilarity As(V) reduction (36) was not significantly different between high vs low As paddy soils (Fig. 3). Because ArrA is induced by As(V), it is possible that the dominant As species, i.e. As(III) in paddy soils (18, 19) did not significantly upregulated its transcriptional activity directly. Moreover, our study showed the increased transcriptional activity of As metabolism genes was due to several distinct clades (community-wide) in the high As paddy soils (Fig. 4 and Supplementary Fig. S4-S12), and this result again corroborated the pervasive presence of As metabolism mechanisms in microbes of the paddy ecosystem.

It’s also noticeable that the relative abundance of *aioA* genes (DNA level) were about 3.5 times higher in high than low As paddy soils, although not statistically significant (*p* = 0.114). In contrast, the relative abundance of *ars* genes at the DNA level was comparable between high and low As levels (Fig. 3A). Considering that the As respiratory As(III) oxidation catalyzed by Aio in microbes can couple As(III) oxidation to ATP production, while the Ars is only responsible for As resistance (14, 34), our finding is consistent with the expectation that energy-production genes (e.g., *aioA*) will be at higher demand at both the DNA and transcriptional (RNA) levels with increased availability of the substrate, while for the resistance genes (e.g., *ars*) transcriptional response might be adequate with increased substrate and the substrate might not be similarly toxic to all species in order to select for the corresponding genes.

The As-associated microbial populations in high As soils mostly clustered together and separate from their low As counterparts based on the similarities among the expression levels of the *ars*, *aioA* and *arrA* genes (Fig. S14), except for sample SKS_AsHig, which could be attributed to high soil heterogeneity or the non-significant effect of As levels on *arrA* genes. In contrast, the As-associated populations did not show any obvious clustering in terms of high As vs low As soils based on metagenomic (DNA) abundances (Fig. S14). These results are overall consistent with the relative abundance and expression of As metabolism genes, indicating that high As select more for transcriptome response as opposed to gene abundance at DNA level.

In terms of whole microbial communities, geographic distances, rather than As levels, were found to shape microbial community differences among the paddy soil sites studied here (Fig. 5). The geographic patterns of soil microbial communities have been widely studied previously and are mostly attributed to varied physicochemical and carbon sources in soil environments (38–40). It is possible that the As levels in these paddy soils did not possess a strong-enough selective pressure on the whole microbial communities. In contrast to As, the concentrations of TP (0.31-0.46 g kg^-1^) and Fe (14.8-27.0 mg kg^-1^), which were about 1.5-1.8 times different between high and low paddy soils, were significant in determining microbial community diversity among our samples (Fig. 5). Previous studies have shown that available organic soil P has an important influence on microbial community composition, shifting both the composition and function of soils microbes (41, 42). Especially for rhizosphere-associated microbes such as the dominant Proteobacteria in paddy soils (43) their community structure can be altered by soil P availability (44, 45). The Fe in paddy soils is likely to impact the microbially mediated redox processes (46), and its presence in the wetland environments has been demonstrated to correlate with microbial biogeographic patterns. It is also important to mention that the effect of Fe and TP on As microbes and genes may be interlinked due to the significant (*p* < 0.01) interaction of P and Fe since P is often the limiting nutrient in many terrestrial ecosystems due to strong sorption on Fe(III) hydroxides (47). A significant positive correlation between P and Fe in porewaters as well as soil matrix has been previously observed in flooded soils with rice growth (48). Moreover, their interaction could also affect As behavior in paddy soils (49) because iron oxyhydroxides could sequester As(V) (50) and reduce As bioavailability to microbes.

Overall, our study revealed that the high As level of ∼100 kg mg^-1^ increased As metabolism genes activities rather than their abundance or diversity in paddy soils. These findings advance our understanding of how microbes respond to high As levels and the diversity of As metabolism genes in paddy soils. Therefore, the potential functional contributions of the ubiquitous As metabolism genes in the paddy ecosystem, and likely other environments, should be examined not only at the DNA level but also at the transcript and possibly protein levels. The As-associated microbial populations in soils with As could have significant contributions to the biogeochemical cycling of As and the MAGs reported here could facilitate future studies in this area. The work presented here could also be served as a guide for the number of samples to obtain, amount of sequencing to apply, and what bioinformatics analyses to perform for studying the dynamics of the As or other elements metabolism pathways.

## MATERIALS AND METHODS

### Sites description and sample collection

Soil samples were collected from six distinct paddy fields in Hunan province, China, including three sites with high As levels (67.3-104.0 mg kg^-1^) in Cili (CL_AsHig), Chenzhou (CZ_AsHig) and Shuikoushan (SKS_AsHig), and three sites with low As levels (2.5-10.8 mg kg^-1^) in Hanpu (HP_AsLow), Lianhua (LH_AsLow) and Yuchangping (YCP_AsLow). Eighteen soil samples, three replicates from the same field for each site, were collected from the soil surface (0-20 cm) in July 2016 during the rice-growing period (flooded). Paddy soil samples were placed in sterile plastic bags and transported to the laboratory on ice for DNA and RNA extraction and soil physiochemistry analysis. Soil properties, including pH, Eh, total concentration of carbon (TC), nitrogen (TN), phosphorus (TP), sulfur (TS), arsenic (As), lead (Pb), cadmium (Cd), chromium (Cr), antimony (Sb) and iron (Fe) were determined on a composite sample of the three replicates following standard methods (21). Details of the underlying methods are provided in the supplementary.

#### Soil DNA and RNA isolation, library preparation and sequencing

Soil DNA and total RNA was extracted from 0.5 g and 2 g of soil using the FastDNA SPIN kit (MP Biomedicals) and E.Z.N.A Soil RNA Midi kit (Omega, Bio-Tek Inc., USA), respectively, according to the manufacturer’s protocol. For each site, DNA or total RNA extracted from the triplicates were pooled together to obtain enough high-quality material for each site and to overcome high sample-to-sample heterogeneity, which is characteristic of soil microbial communities. The quality of DNA and total RNA was assessed by NanoDrop ND-1000 spectrophotometer (Thermo Fisher Scientific, Inc., Wilmington, DE, USA) and the ratio of absorbance at 260 nm and 280 nm was around 1.8-2.0, indicating good quality for downstream analysis. Total RNA of the paddy soils was subjected to an rRNA removal procedure using the Ribo-Zero rRNA removal bacteria kit (Illumina Inc., San Diego, CA, USA) following the manufacturer’s protocol.

Paired-end DNA and cDNA libraries (2 × 150 bp) were constructed using TruSeq^TM^ DNA Sample prep kit (Illumina Inc., San Diego, CA, USA) and TruSeq^TM^ RNA Sample prep kit (Illumina Inc., San Diego, CA, USA), respectively, following the manufacturer’s protocol. The six DNA and cDNA libraries were sequenced respectively on an Illumina HiSeq 3000 platform at Majorbio Bio-pharm Technology Company in Shanghai, China with HiSeq 3000/4000 PE Cluster kit and HiSeq 3000/4000 SBS kit (Illumina Inc., San Diego, CA, USA) according to the manufacturer’s protocol.

#### Taxonomic classification, functional annotation, and population genome binning of soil metagenomes and metatranscriptomes

The raw paired-end reads of metagenomic and metatranscriptomics datasets were trimmed using SolexaQA (51), removing reads with Phred quality score (Q < 20) and minimum fragment read length of 50 bp after trimming for downstream analyses. Parallel-META version 2.4 (52) was used with default settings to recover 16S rRNA gene fragments from metagenomes and to identify and remove residual rRNA sequences after rRNA subtraction from metatranscriptomes. 16S rRNA gene fragments from metagenomes were then processed for Operational Taxonomic Unit (OTU) picking using close OTU picking, defined at the 97% nucleotide sequence identity threshold, and taxonomic identification with the SILVA database (29) as implemented in QIIME 1.9.1 (53) and classified at the phylum and class levels. The relative abundance of each taxon at each level was normalized by the total number of 16S rRNA gene sequence fragments recovered from the corresponding metagenome.

Protein-coding sequences from short-read metagenomes and metatranscriptomes were predicted using FragGeneScan (54) with default settings and searched against Swiss-Prot database (UniProt, downloaded in April 2019) using BLASTp (BLAST + version 2.2.28) (55). Only the best match with amino acid identity ≥ 50% and query sequence length coverage ≥ 70% by the alignment was retained. Protein-coding sequences were also functionally annotated, using the same threshold, against the SEED database using the subsystem categories (56). Differentially abundant taxa based on 16S rRNA gene sequence fragments and SEED subsystems of metagenomics and metatranscriptomics between high and low As levels were determined with the DESeq2 package using the negative binomial model, adjusted for false discovery rate (57). Heatmap in pheatmap R package (R version 3.5.1) (58) was used to visualize the significantly differentially abundant taxonomic levels and SEED subsystems (adjusted *p* value < 0.05). Metagenome-assembled genomes (MAGs) were recovered using both MaxBin Version 2.1.1 (59) and MetaBAT Version 2.12.1 (60) as described in more detail in supplementary.

#### Assessment of the abundance and transcriptional activity of arsenic metabolism genes

##### Building an As metabolism protein reference database

To identify and quantify As metabolism proteins containing reads, in-house databases of eight As resistance proteins [Ars i.e. ArsR (ArsR1, ArsR2, ArsR3 and ArsR4), ACR3, ArsB, ArsC (ArsC1, ArsC2 and ACR2), ArsM, ArsI, ArsP, ArsH] and As respiratory oxidation (AioA) and reduction (ArrA) proteins were constructed and manually curated as described before (5). Briefly, verified arsenic metabolism protein sequences were downloaded from UniProt or NCBI’s NR database to build HMM profiles, and searched against 786 representative species which are sampled from an original dataset used for reconstruction of tree of life (species tree) (61), to identify homologs of As metabolism proteins. The resulting matching sequences were manually inspected for the presence of conserved domains based on the Pfam models (release 28.0) (62) and forming deep-branching clades in the phylogenetic tree of the (manually) verified reference sequences to further determine if the sequence in question was truly As metabolism protein. The tree was built using maximum likelihood as implemented in RAxML v8.4.1 with the PROTGAMMAAUTO model (63). Sequences forming long-branches and/or lacking one or more of the conserved domains were removed from further analysis.

To build a more comprehensive As metabolism proteins database that better captures the diversity of As metabolism proteins, proteins on the contigs assembled using IDBA with default settings (64) from metagenomes and metatranscriptomes (for more details, see supplementary) were also searched against the curated As metabolism proteins database using BLASTp (BLAST + version 2.2.28) with a cut-off for a match of amino acid identity ≥ 40% and reference length coverage ≥ 70% by the alignment. A total of 1040 Ars proteins, including 87 ACR3, 281 ArsC, 1 ArsH, 69 ArsI, 173 ArsM, 21 ArsP and 408 ArsR were identified using this approach. However, for the AioA and ArrA proteins that are longer than the Ars proteins (800-900 vs 200-400 amino acids), no match was identified, and it was likely attributable to the limiting assembly, consistent with the relatively low sequencing effort. Therefore, in order to obtain a comprehensive database for AioA and ArrA proteins, the AioA and ArrA references in curated As metabolism database were searched against UniRef90 (TrEMBL, downloaded in October 2018) database for additional homologs. A more stringent cut-off compared to the Ars proteins mentioned above, i.e. identity ≥ 40% and reference length coverage ≥ 90% by the alignment, was used to filter the AioA and ArrA proteins in UniRef90 database. Identified Ars protein sequences from metagenomes and metatranscriptomes, and AioA and ArrA proteins sequences from UniRef90 database were searched against Pfam database (release 32.0) (62) in Jun 2019 and inspected for the presence of known conserved domains. The resulting matching sequences were aligned using MAFFT version 7.407 (65) and visually inspected for forming deep-branching clades in a maximum likelihood phylogenetic tree as described above to determine if the sequences were truly As metabolism proteins. These phylogenetic trees were also used for placement of As metabolism proteins encoding reads as described below.

##### Estimation of As metabolism genes abundance and transcriptional activity

Short sequences from metagenomes and metatranscriptomes were searched against the comprehensive Ars, AioA and ArrA references databases using BLASTx search (BLAST + version 2.2.28) and filtered for best matches with amino acid identity ≥ 90%, alignment length ≥ 25 amino acid and e-value ≤ 1E-5. MicrobeCensus was used to assess the average genome size of each sample community (66) and normalize the total base-pairs assign to each As gene by gene length, i.e. RPKM = (the number of reads mapping on reference sequences/gene length in kb per million reads), and genome equivalent, i.e., RPKM/average genome size, providing an estimate RPKG [(reads mapped to gene)/(gene length in kb)/(genome equivalents)] of how many cells from those sampled encode the gene of interest. Relative abundance of As metabolism genes in metatranscriptomics was calculated by the statistic RPKM (reads per kb per millions of reads), i.e. RPKM = (the number of reads mapping on reference sequences)/(gene length in kb)/(the total number of reads after removal of rRNA reads). The As metabolism genes transcriptional activity was evaluated by the relative abundance of As metabolism genes in metatranscriptomes (RPKM) normalized by the relative abundance of the same gene variant or OTU (at > 90% amino acid level) identified in metagenomes (RPKM; we used RPKM to normalized the relative abundance of the same gene identified in metagenomes here instead of RPKG in order to keep the normalization criteria consistent between metagenomes and metatranscriptomes in calculating the gene’s transcriptional activity), and visualized in heatmap using the pheatmap package in R 3.5.1 (58). One-way ANOVA analysis, performed with the vegan package (67) in R 3.5.1, was used to compare the difference of As metabolism genes abundances and transcriptional activities in high vs low As levels paddy soils.

##### Phylogenetic placement of As metabolism gene carrying reads

Arsenic metabolism protein-containing reads identified by BLASTx search were firstly translated to amino-acid sequences by FragGeneScan (54) with default settings. The amino acid fragments were then added to the As metabolism protein references alignments using MAFFT version 7.407 (65) with ‘addfragments’ option and were placed in the corresponding phylogenetic tree of different As metabolism proteins (described above) respectively, using RAxML v8.4.1 (63) with -f v option. The placement of the target As metabolism protein-containing reads was visualized in iTOL (68) after processing the resulting visualization jplace file generated by RAxML with JPlace.to_iTOL.rb from the Enveomics Collection (69).

#### Alpha and beta community diversity estimation

The abundance-weighted average coverage of the sampled microbial communities achieved by our sequencing effort was estimated by Nonpareil (70, 71) based on the level of redundancy of the sequence reads of metagenomes. Mash distances (72) were used to assess the sample-to-sample sequence composition similarity at the whole metagenome level, based on shared kmers. Because the coverage of these metagenomics varied among different studies, the datasets were subsample to 2Gbp to make them comparable. Principle coordinates analysis (PCoA) and/or non-metrical multidimensional scaling (NMDS) plot were used to visualize the site-to-site similarity matrices of microbial community compositions based on Mash distances, OTU tables, and annotation counts from SEED subsystems based on Bray-Curtis distance. NMDS plot of annotation counts of *ars* [i.e. *arsR* (*arsR1*, *arsR2*, *arsR3*, *arsR4*), *acr3*, *arsB*, *arsC* (*arsC1*, *arsC2* and *acr2*), *arsM*, *arsI*, *arsP*, *arsH*], *aioA*, *arrA* genes from metagenomes and metatranscriptomes based on Bray-Curtis distances were used to visualize the site-to-site similarity of As-associated microbial community compositions.

The Adonis test was used to determine correlations between physicochemical variables measured at each site and similarity in microbial community composition (Mash distance) or *ars* genes abundances/transcriptional activities (Bray-Curtis distances) and tested for significance based on a permutation test with 999 interactions. The interactions between different physicochemical variables were assessed by PERMANOVA analysis while the correlation of physicochemical variables with *aioA* and *arrA* gene abundances and transcriptional activities in different sites were tested by Pearson correlation. These analyses were performed in R 3.5.1 with the vegan (67) and ggplot2 (73) packages.

#### Data availability

Metagenomics and metatranscriptomics datasets have been deposited in National Center for Biotechnology Information (NCBI)’s Short Read Archive (SRA) database, under the bioproject PRJNA616041. NCBI SRA numbers for all datasets were provided in supplementary Table S1.

## Acknowledgements

This research was supported, in part, by the China Postdoctoral Science Foundation (No. 212400241) and Major Project of the Ministry of Science and Technology of Jiangxi Province (No. CK201302055). KTK’s research was supported, in part, by the U.S. National Science Foundation (Award No. 1831582 and 1759831). We also appreciate the support from the Brook Byers Institute for Sustainable Systems and the Hightower Chair at the Georgia Institute of Technology.

